# Pseudotime analysis for time-series single-cell sequencing and imaging data

**DOI:** 10.1101/2023.11.03.565575

**Authors:** Gang Li, Hyeon-Jin Kim, Sriram Pendyala, Ran Zhang, Christine M. Disteche, Jean-Philippe Vert, Xinxian Deng, Douglas M. Fowler, William Stafford Noble

## Abstract

Many single-cell RNA-sequencing studies have collected time-series data to investigate transcriptional changes concerning various notions of biological time, such as cell differentiation, embryonic development, and response to stimulus. Accordingly, several unsupervised and supervised computational methods have been developed to construct single-cell pseudotime embeddings for extracting the temporal order of transcriptional cell states from these time-series scRNA-seq datasets. However, existing methods, such as psupertime, suffer from low predictive accuracy, and this problem becomes even worse when we try to generalize to other data types such as scATAC-seq or microscopy images. To address this problem, we propose Sceptic, a support vector machine model for supervised pseudotime analysis. Whereas psupertime employs a single joint regression model, Sceptic simultaneously trains multiple classifiers with separate score functions for each time point and also allows for non-linear kernel functions. Sceptic first generates a probability vector for each cell and then aims to predict chronological age via conditional expectation. We demonstrate that Sceptic achieves significantly improved prediction power (accuracy improved by 1.4 *−* 38.9%) for six publicly available scRNA-seq data sets over state-of-the-art methods, and that Sceptic also works well for single-nucleus image data. Moreover, we observe that the pseudotimes assigned by Sceptic show stronger correlations with nuclear morphology than the observed times, suggesting that these pseudotimes accurately capture the heterogeneity of nuclei derived from a single time point and thus provide more informative time labels than the observed times. Finally, we show that Sceptic accurately captures sex-specific differentiation timing from both scATAC-seq and scRNA-seq data.

## 1 Introduction

With the decreasing cost of single-cell RNA-sequencing (scRNA-seq), it is increasingly common to collect time-series scRNA-seq data sets. Such data can be used to investigate transcriptional changes over various notions of biological time, such as cell differentiation [1], embryonic development [13] and response to stimulus [21].

However, analysis of any time series single-cell dataset requires that we distinguish between “time” and “pseudotime.” The first of these terms is well-defined: “time” simply refers to the time point at which each sample was collected. “Pseudotime,” on the other hand, refers to a “quantitative measure of progress through a biological process” [20]. The term was introduced in the context of single-cell genomics as a way to segregate a collection of measured cells along a developmental trajectory. The key idea is that measurement of a heterogeneous population of cells collected at a single time point may still capture cells in many different stages of development.

Accordingly, many pseudotime inference procedures have been developed, each attempting to automatically infer pseudotimes for individual cells, frequently with respect to branching trajectories. Principal components analysis (PCA) [11] is commonly regarded as a straightforward and interpretable baseline method for pseudotime inference [10]. Slingshot identifies cell lineages by treating groups of cells as nodes within a graph and identifying a minimum spanning tree connecting these nodes [18]. Monocle 2 [14] and Monocle 3 [4], which have been widely adopted [15], employ distinct underlying algorithms. Monocle 2 utilizes a reversed graph embedding strategy to model cell trajectories, effectively constructing a minimum spanning tree among cells [14]. Monocle 3 identifies cell trajectories using a single-rooted directed acyclic graph, which captures the hierarchical organization of cell states [4]. Finally, Palantir is designed to model the trajectories of differentiating cells by treating the determination of cell fate as a probabilistic process, using entropy as a key metric to quantify cell plasticity as cells progress [17]. Each of these methods is typically applied to data derived from a set of cells collected at a single time point.

When single-cell data is collected from multiple time points, we must somehow integrate these two notions of time. At least four different methods for achieving this integration are possible. First, some early time series scRNA-seq studies ignored pseudotime entirely, opting to focus on how gene expression changes as a function of time [6]. Second, one can ignore the time labels by applying a pseudotime inference algorithm to the concatenated dataset. This is the most commonly adopted approach. Third, in principle one could run a pseudotime inference algorithm separately on each time point and then, in a post-processing step, somehow calibrate the pseudotime labels across time points. However, we are not aware of any methods that have attempted this type of *post hoc* calibration.

The fourth approach for integration of time and pseudotime—inferring pseudotimes while taking into account the observed time labels—is the one we pursue here. Several previous methods have adopted related approaches. For example, Tempora focuses on delineating how cell types are interrelated throughout a time series dataset [19]. The method first clusters cells into nominal cell types and infers a trajectory at the cell type level. The directionality of each edge in this cell-type trajectory is then inferred using the associated time labels. Subsequently, Psupertime proposed transforming the unsupervised learning problem of assigning pseudotimes to cells into a supervised problem, by using the observed times as labels in a regression setup [10]. Their psupertime software uses a penalized ordinal logistical regression model for time-series scRNAseq data As in several other cases where supervised methods have been used to improve upon unsupervised single-cell analyses [23, 24], psupertime has been shown to perform comparably to or better than competing methods, including PCA, Monocle 2, Slingshot and Tempora.

We hypothesized that psupertime’s predictive accuracy suffers from its use of a simple linear combination of gene expression profiles. Motivated by this hypothesis, we developed Sceptic, a support vector machine (SVM) framework for supervised pseudotime analysis. Sceptic trains a series of onevs-the-rest classifiers, thereby generating for each cell a probability vector over all the time points in the dataset. Sceptic then predicts each cell’s pseudotime via conditional expectation. We validated the performance of Sceptic on multiple public single-cell datasets and further extended it to different modalities of time-series single-cell data, including our own single-nucleus imaging dataset. First, we demonstrated that Sceptic achieves higher accuracy than psupertime on six scRNA-seq datasets. Second, we showed that Sceptic works well for single-nucleus imaging data. Third, we applied Sceptic to scATAC-seq data and showed that the model captures sex-specific differentiation from both scATAC-seq and scRNA-seq data. Finally, we applied Sceptic to a co-assay data set, where we detected a methylation delay that agrees with other independent studies. Sceptic’s Python code can be found, with an Apache license, at https://github.com/Noble-Lab/Sceptic.

## 2 Methods

### 2.1 Single CEll PseudoTIme Classifier (Sceptic)

Sceptic is an SVM-based model for pseudotime analysis. It takes as input a single-cell measurement matrix *X*, with *M* rows of measurements and *N* columns corresponding to cells, and a corresponding time label vector 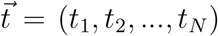. We assume that the cells are collected from *D* different observed time points, i.e., *t*_*i*_ *∈ {T*_1_, *T*_2_, …, *T*_*D*_*}*. Sceptic trains one support vector machine classifier for each timestamp, say *T*_1_, optimizing a coefficient vector **a** = (*a*_1_, *a*_2_, …, *a*_*N*_) for *T*_1_ to minimize the following loss with 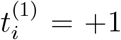 if *t*_*i*_ = *T*_1_ and 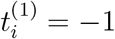 if *t*_≠_*T*_1_:

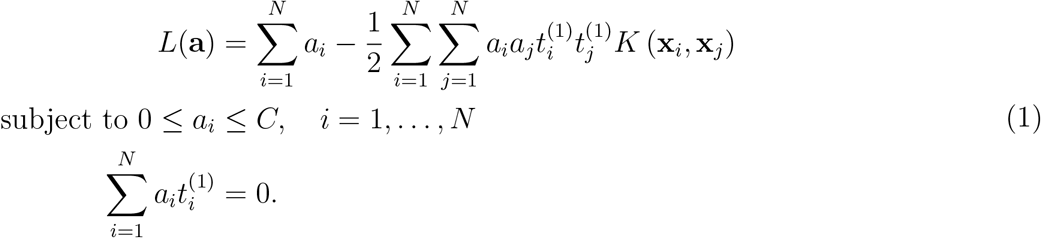

where *K* (*x*_*i*_, *x*_*j*_) = *ϕ* (*x*_*i*_)^*T*^ *ϕ* (*x*_*j*_) is the kernel function. Sceptic default uses a radial basis function (RBF) kernel *K* (*x*_*i*_, *x*_*j*_) = exp *−*γ ∥*x*_*i*_ *− x*_*j*_∥^2^ but also allows other kernels such as a linear kernel 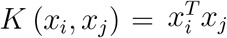. Once fitted, Sceptic estimates a class membership probability **P** = (*p*_1_, *p*_2_, …, *p*_*D*_) where p_*d*_ is the probability that the cell is collected from time point *T*_*d*_ based on Wu *et al*. [22]. The pairwise estimated probabilities between any pair of classes are first calibrated using Platt scaling and then used to solve a quadratic optimization where the solution is the class membership probability. Finally, the predicted pesudotime *Y* for cell *x* is given by

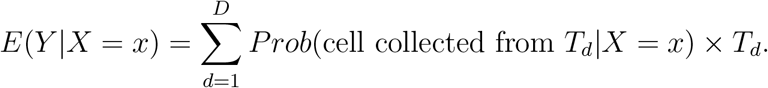

#### Nested cross validation

To select hyperparameters while avoiding overfitting, we train Sceptic using a nested cross-validation scheme. Cross-validation involves randomly splitting the data into, say, five equal-sized folds, and then iteratively using four folds to train the model and the remaining fold for testing. Nested cross-validation repeats another cross-validation splitting procedure on each training set. We use the internal 4-fold cross-validation step to select a set of optimal hyperparameters for each training set, and these hyperparameters are then used to train a model on the full training set. The trained model is then used to make predictions on the test set. We repeat this process five times as the external 5-fold cross-validation step to make predictions on all the data.

#### Hyperparameter search

In the inner loop of the nested cross-validation, Sceptic performs a grid search over three hyperparameters. The first hyperparameter *R* is a Boolean indicating whether the SVM uses a linear or RBF kernel. The second hyperparameter *C* controls the relative weight assigned to violations of the SVM soft margin. The third hyperparameter γ specifies the inverse of the width of the RBF kernel. We adopt sklearn’s default γ for support vector classifier: γ = 1/(*M * Var*(*vector*(*X*)), which adjusts the number of features *M* and the variability of the measurement matrix via the variance of the measurement matrix *X* in vector format. Sceptic employs a six-valued grid with dimensions *R ∈ {*0, 1*}, C ∈ {*0.1, 1, 10*}*.

### 2.2 Psupertime

For five datasets that psupertime evaluated, we directly compared Sceptic’s results with the reported performance in the psupertime paper. For mESC data, we follow their tutorial using the default setting to select the hyperparameter *λ* for the weight of the L1 regularization penalty, selecting the *λ*.1se with performance within one standard error of the best performance *λ*.min, and then re-training the model using all the cells.

### 2.3 Simulation

We generated a random set of 200 cells, each assigned to one of five observed time points: day 0, 3, 7, 11, and 21. For each cell, the simulated pseudotime was obtained by sampling from a Gaussian distribution with a mean corresponding to its observed time point. Then the scale of the pseudotime was normalized to a range between 0 and 21, while preserving the ordering of the cells along the pseudotime continuum. This normalization ensures consistency in the pseudotime scale while maintaining the temporal order of the cells.

We first simulated a linear differentiation process. To do so, we performed a linear projection of the normalized pseudotime onto a 500-dimensional gene expression space. This projection was achieved by multiplying the normalized pseudotime values by 500 random Gaussian projection parameters. The result is a 200-by-500 matrix representing the cell-by-gene expression, where each row corresponds to a cell, and each column corresponds to a gene. The dataset is associated with 200 observed time points, and 200 normalized pseudotime values need to be estimated.

Next, we simulated a bifurcating differentiation process. In this setting, for cells corresponding to days 7, 11, and 21, each cell is randomly assigned to one of two branches with a 0.5 probability. Subsequently, the normalized pseudotime for each branch is projected onto a 500-gene expression space by multiplying it with 500 random Gaussian projection parameters specific to that branch. The cubic power transformation is then applied to all genes to simulate the non-linear relationship between genes and pseudotime. This process results in a 200-by-500 matrix representing cell-by-gene expression. As in the first simulation, the dataset is associated with 200 observed time points, and 200 normalized pseudotime values need to be estimated.

Descriptions of the scRNA-seq, scATAC-seq, and imaging datasets are provided in Supplementary Section 1.

## 3 Results

### 3.1 Overview of Sceptic, Single CEll PseudoTIme Classifier

Sceptic can perform pseudotime analysis on various types of single-cell or single-nucleus data. The model takes as input a collection of such data, learns the relationship between the observed data and the associated time stamps, and finally uses the trained model to assign to each cell a real-valued pseudotime. Ideally, the pseudotimes assigned by Sceptic reflect each cell’s progression along a notion of time—developmental, cell cycle, disease progression, aging—that is appropriate to the given data.

Sceptic differs from its predecessor, psupertime [10], in three ways (Figure 1). First, Sceptic uses a nonlinear support vector machine rather than an ordinal logistic regression model. Second, and more importantly, whereas psupertime trains a single regressor with multiple thresholds, Sceptic trains a collection of classifiers, one for each timestamp, using a one-versus-the-rest strategy. The final pseudotime assigned to a given cell is computed as a weighted sum of the different predictions. This setup significantly enhances the classifier’s performance. Third, Sceptic employs a cross-validation strategy, in which a collection of models is trained on different subsets of the data and used to make predictions on disjoint test sets. This strategy prevents the model from reporting pseudotime values that may overfit the data.

**Figure 1:**
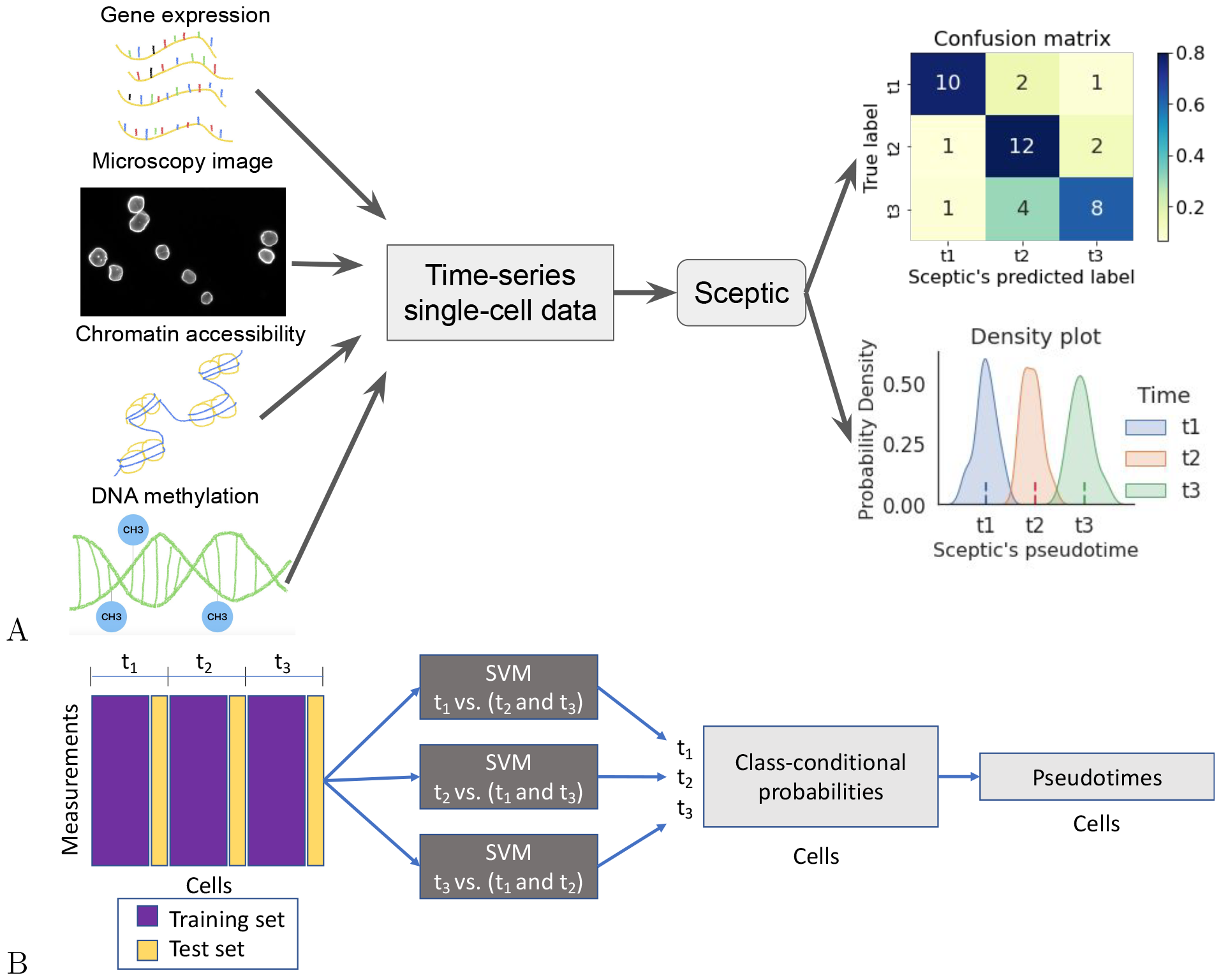
Sceptic analysis for time-series single-cell sequencing and imaging data. **A** Sceptic from the user’s perspective. Sceptic can take various types of single-cell or single-nucleus data as input and is trained to predict each cell’s time label (shown as a confusion matrix) and to infer the pseudotime for each cell (shown as a density plot across all cells). **B** Overview of Sceptic analysis.

As motivation, we created two simple simulations to illustrate how Sceptic works (details in Section 2.3). The simulation framework is modeled after an existing collection of mouse embryonic stem cell (mESC) data [1], and the simulation approach is inspired by the one employed by dynverse [15]. We simulated two scenarios: linear differentiation and a bifurcating structure. In each scenario, we compared Sceptic with psupertime and a ridge regression baseline. The results (Figure 2) suggest that, in the linear differentiation setting, all three methods accurately preserve the simulated ordering of cells; however, only Sceptic and ridge regression accurately predict the true pseudotime values. Psupertime’s predictions, in contrast, are a monotonic transformation of the true pseudotime. In the bifurcating structure setting, Sceptic achieves the best prediction among all three methods by not only preserving the ordering of cells but also predicting the correct scaling of the pseudotimes. Psupertime fails to predict the correct ordering of cells, and ridge regression’s predictions do not reflect the actual scale of the simulated pseudotimes.

**Figure 2:**
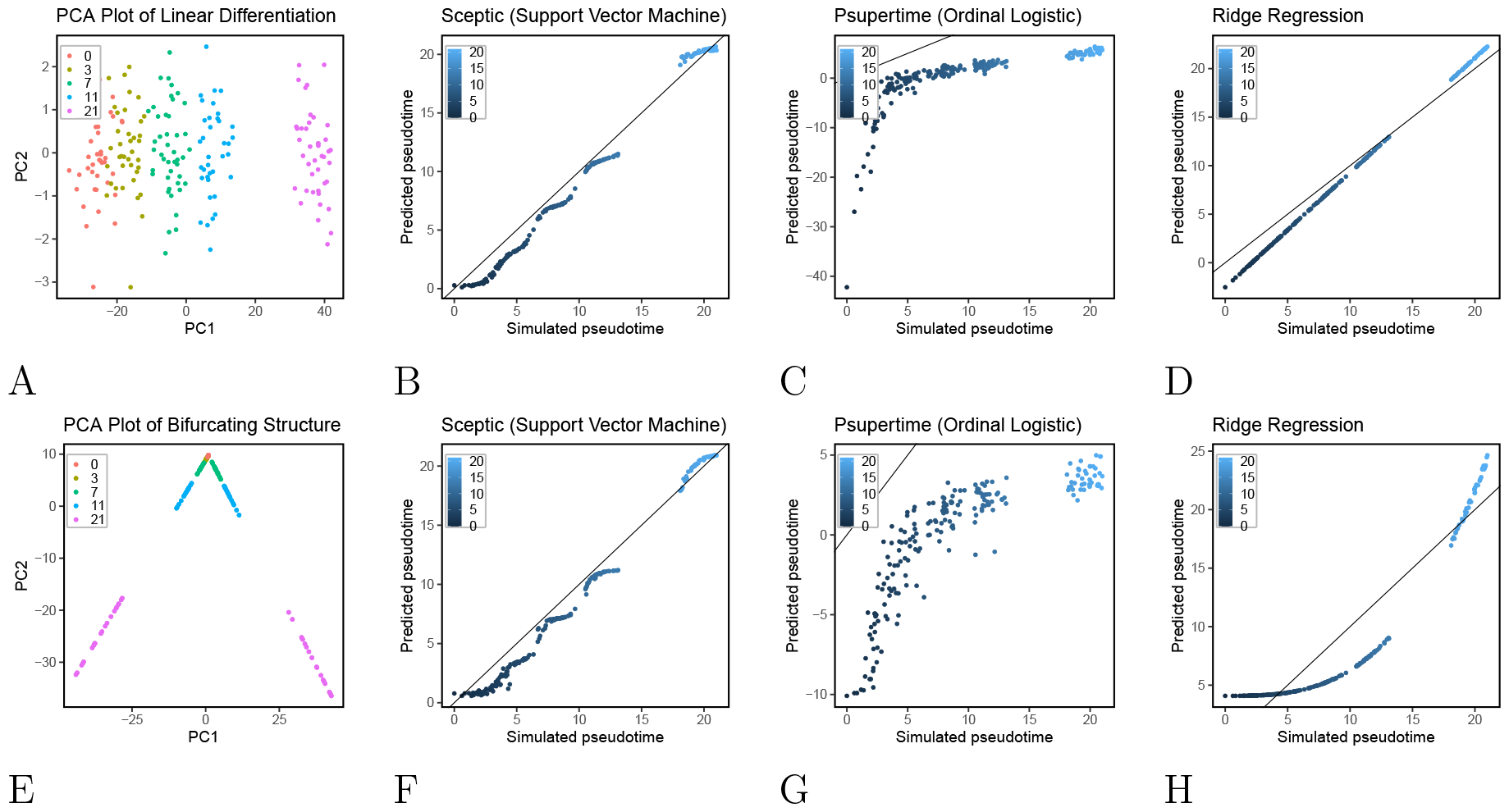
Sceptic works well in simulation. **A** PCA visualization of linear differentiation data. **B** Sceptic’s performance for linear differentiation data. **C** Psupertime’s performance for linear differentiation data. **D** Ridge regression’s performance for linear differentiation data. **E** PCA visualization of bifurcating structure data. **F** Sceptic’s performance for bifurcating structure data. **G** Puspertime’s performance for bifurcating structure data. **H** Ridge regression’s performance for bifurcating structure data.

### 3.2 Sceptic outperforms psupertime on classification for time-series singlecell RNA-sequencing data

We designed Sceptic to provide a flexible framework for modeling pseudotime trends within a collection of single-cell data. Accordingly, we set out to test the hypothesis that Sceptic could more accurately predict the timestamp associated with a given cell than psupertime. To do this, we used our recently published scRNA-seq data from a mouse embryonic stem cell (mESC) differentiating time-series dataset, collected over five time points: days 0, 3, 7, 11 and neural progenitor cells collected at day 21 [1]. We used five-fold cross-validation, training a model on 80% of the cells and testing on the remaining 20%, and repeating this procedure five times. For comparison, we also applied psupertime to the same dataset. Because psupertime is not designed to generalize to new cells, we are only able to report psupertime’s performance when fitting a single model to the entire dataset. In this sense, the comparison is biased in favor of psupertime.

Despite this bias, the predictions from the two models, summarized in confusion matrices (Figure 3A), show that Sceptic outperforms psupertime. In particular, Sceptic achieves an accuracy of 93.73% (because 3,809 of the 4,064 predictions fall along the diagonal of the confusion matrix), whereas psupertime’s accuracy is only 89.94% (3,655 correct predictions). This difference is significant (*p* = 4.94*e*^−10^) according to Fisher’s exact test. The confusion matrices indicate which timestamps are difficult to differentiate, with most of the off-diagonal predictions occurring between days 3 and 7 and between days 7 and 11. Sceptic achieves a clear separation between cells of day 7 and 11 (Figure 3B), whereas psupertime does not perform very well on these cells (Figure 3A).

**Figure 3:**
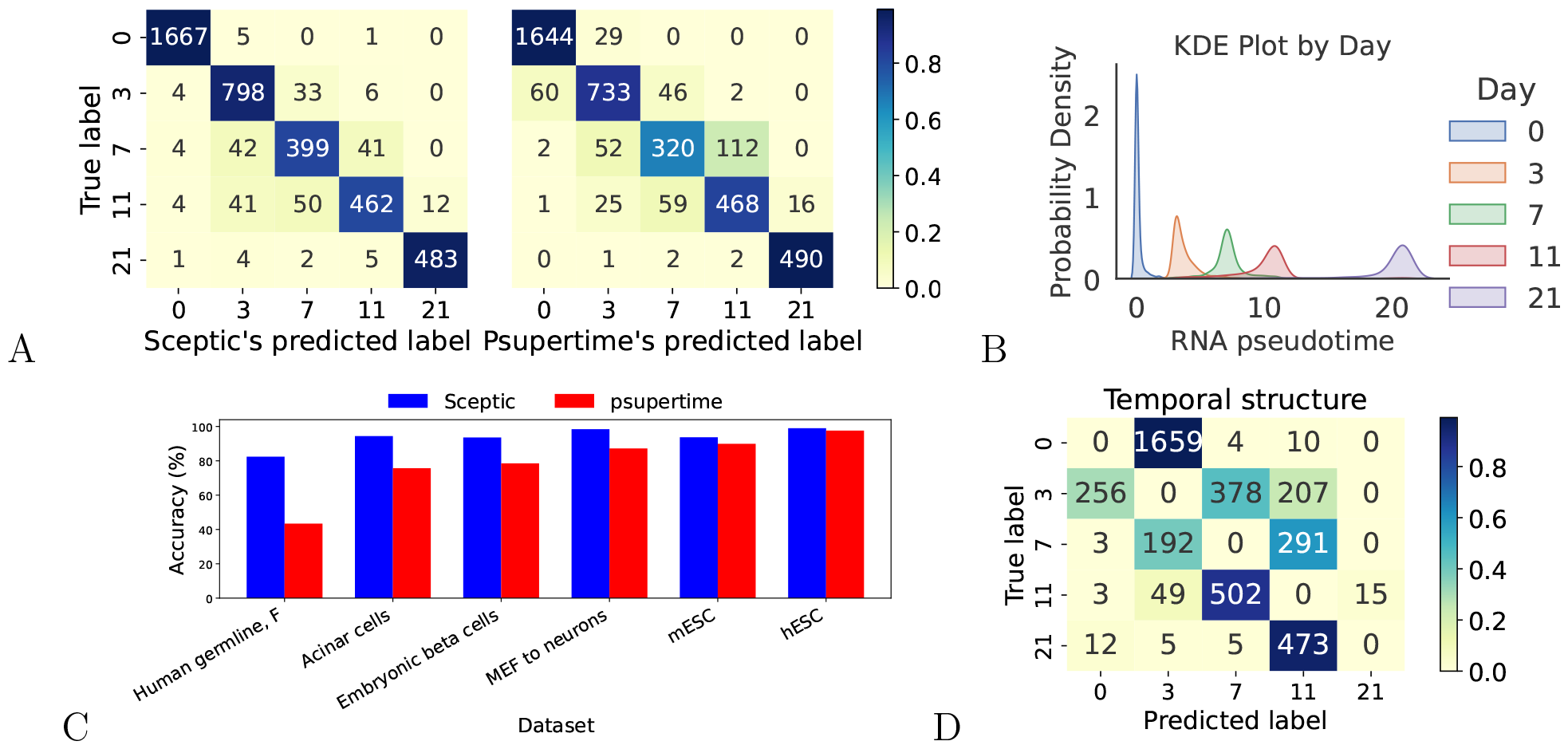
Sceptic achieves better classification accuracy than psupertime on single-cell RNA sequencing data. **A** Confusion matrices for Sceptic (left) and psupertime (right) on the mESC data. Sceptic results use cross-validation, whereas psupertime results are from training on the full dataset. **B** Kernel density plot of Sceptic pseudotime, colored by the day of cells, on the mESC data. **C** Accuracy of both methods on six time-series datasets. **D** Confusion matrix from leave-one-time-point-out analysis. A successful model would tend to predict time points adjacent to the diagonal.

To further test our hypothesis, we downloaded the five datasets that were analyzed in the original psupertime paper [10], and we repeated the above analysis. In this case, for psupertime we use the previously reported accuracy values. Again, despite the bias in favor of psupertime in this setup, on all five datasets Sceptic achieves consistently higher classification accuracy than psupertime (Figure 3C), improving by 1.4%-39.0%. The difference is significant (*p* = 0.03125) according to a Wilcoxon signed-rank test. Note that psupertime has previously been shown to perform comparably to or better than other methods like PCA [11], Monocle 2 [14], Slingshot [18] and Tempora [19]. Thus, we only compare our performance with psupertime.

We note that, in any evaluation of a supervised pseudotime inference method, we face the fundamental challenge of distinguishing between batch effects and true pseudotemporal ordering. To understand the problem, consider a theoretical dataset in which each time point is coupled with a very strong, time-specific batch effect. In such a setting, a method that successfully learns to identify this batch effect will perform perfectly in a confusion matrix, despite not having learned anything about the temporal signal in the data. Therefore, additional validation is required to demonstrate that the pseudotemporal ordering is meaningful.

To address this concern, we repeated our analysis but used a cross-validation strategy in which we iteratively left one time point out of the training set on mESC dataset. In this setting, the resulting confusion matrix (Figure 3D) necessarily has zeroes on the diagonal. Our performance measure is the proportion of values that fall in boxes that are adjacent to the diagonal. The reasoning here is that if the model learned only batch effects, then the errors would be uniformly distributed across all time points. On the other hand, a model that successfully learns temporal patterns will, when forced to predict a time for a novel time point, tend to predict adjacent time points. In our case, we find that 92.7% of the predictions in the leave-one-time-point-out setting fall into adjacent time points. This is substantially greater than the 40.0% that would be expected if the errors were uniformly distributed over time points. Note that generalizing this experiment to a leave-two-time-points-out setting would be challenging, because this datset only has five time points in total.

### 3.3 Sceptic works well for single-nucleus imaging data

Having established that Sceptic’s machine learning framework works well for scRNA-seq data, we next investigated whether a similar approach would generalize to single-nucleus microscopy imaging data. Accordingly, we collected F121-6 mESC differentiation 3D images (DNA staining with DAPI and immunostaining of nucleolin) using the same five time points as described in our recent study [1] (Figure 4A). We segmented each 2D-maximum-projection image using a pre-trained Mask R-CNN model [2, 9], and we used a pre-trained ResNet34 model to generate 1024-dimensional representations for each nucleus (Figure 4B). To measure the performance of the classifier, we randomly selected 1,200 single-nucleus image crops from each time point, yielding 6,000 crops in total.

**Figure 4:**
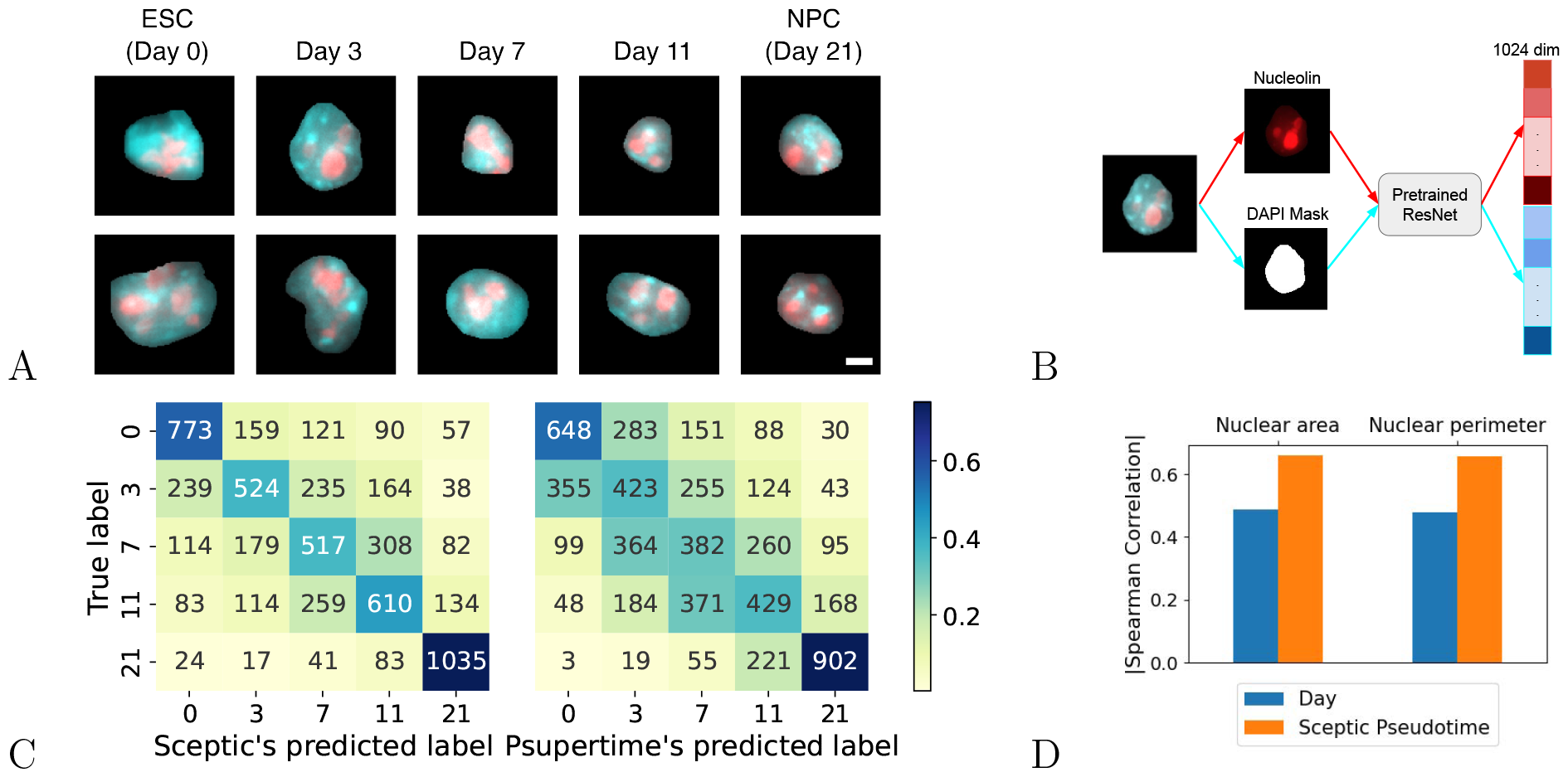
Sceptic works well for single-nucleus imaging data. **A** Examples of single nucleus images. Blue = DAPI; red = nucleolin. Scale bar is 10 *µ*m. ESC: embryonic stem cells. NPC: neural progenitor cells. **B** Image processing pipeline. **C** Sceptic achieves better classification accuracy than psupertime for time-series single-nucleus image data. **D** Pseudotimes inferred by Sceptic are more highly correlated with nuclear morphology features than the observed time label.

Our results show that Sceptic is able to accurately assign time labels to nuclei on the basis of microscopy images, significantly outperforming psupertime on this task (Figure 4C). Overall, Sceptic achieves 57.7% test set accuracy, whereas psupertime achieves a significantly lower 46.4% training set accuracy (*p* = 1.08*e*^−8^), Fisher’s exact test). For both methods, the overall accuracy is lower in the microscopy setting than in the scRNA-seq setting. As expected, Sceptic achieves the best separation between the ends of the time series—day 0 versus day 21.

Next we investigated whether the time labels assigned by Sceptic show stronger correlations with cell morphology features than the observed time labels. During embryonic stem cell differentiation, neural progenitor cells (day 21) exhibit reduced nucleolus activity, reduced transcription of rRNA genes, and smaller nucleoli [8]. Thus, we expect an accurate time label to be strongly correlated with these morphology features. We therefore compared the correlations between pseudotime or observed time with respect to two features: nuclear area, and nuclear perimeter. Compared to the observed time points, Sceptic assigns much finer-grained continuous time labels, and thus achieves a higher absolute correlation with nuclear image morphology features compared with the discrete-time labels (Figure 4D). These results suggest that the pseudotime values inferred by Sceptic accurately capture the heterogeneity of nuclei sampled at the same time point and thus provide more informative time labels than the observed ones.

### 3.4 Sceptic detects sex-specific differentiation from scATAC-seq and scRNAseq data

Having established that Sceptic can work well for both scRNA-seq and single-nucleus imaging data, we next investigated whether a similar approach could generalize to single-cell transposase-accessible chromatin with sequencing (scATAC-seq) data. This assay identifies, in a cell-specific fashion, punctate regions of open chromatin, which typically correspond to various types of regulatory elements, including promoters, enhancers, and insulators. To test Sceptic, we downloaded F121-6 (female) and F123 (male) mESC differentiation scATAC-seq data from Bonora *et al*. [1]. We randomly downsampled the number of cells from each time point to the smallest cell number across all time points, which left us with 535 female cells (107 cells for each time point) and 560 male cells (140 cells for each time point). Note that the F123 male dataset does not contain neural progenitor cells for day 21. We trained two separate Sceptic models for each dataset, one for each sex.

In this experiment, Sceptic achieves 76.8% test set accuracy on F121-6 female cells and 76.1% on F123 male cells (Figure 5A–B). We note that Sceptic accurately classifies cells from day 0 and day 3 for both male and female cells, whereas, on both datasets, cells from days 7 and 11 are difficult to distinguish. Cells from day 7 and day 11 share similar chromatin accessibility patterns, and they construct a single cell trajectory branch (embryonic body branch) in Bonora *et al*. ‘s analyses. In addition, we observed that for day 7, female cells have a slightly higher proportion of cells classified as earlier time points (day 0 and day 3) than their male counterparts (Figure 5A–C): 17.8% (19 out of 107 cells) for female and 9.3% (13 out of 140 cells) for male. Similar differences in proportion are observed for cells in day 11 (34.3% for female cells and 29.0% for male cells). These differences are potentially indicative of female-delayed onset of differentiation and are consistent with previous observations [16, 7, 1], in which the early establishment of X chromosome inactivation is required for differentiation.

**Figure 5:**
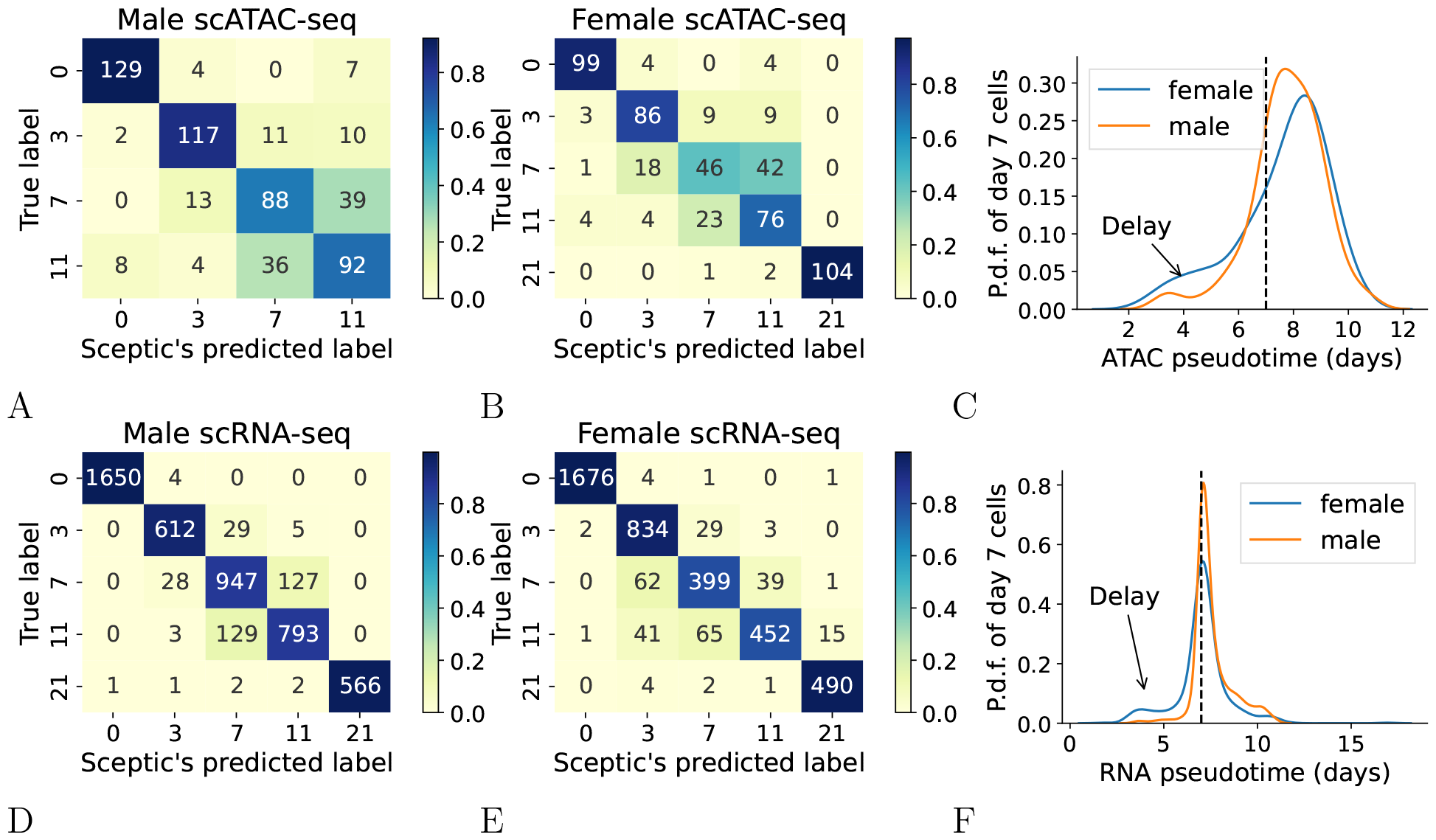
Sceptic applied to scATAC-seq and scRNA-seq mESC data. **A** Sceptic’s performance for scATAC-seq data of F123 (male) cells. **B** Sceptic’s performance for scATAC-seq data of F121-6 (female) cells. **C** Probability density function of Sceptic’s pseudotime for day 7 cells of scATAC-seq data. **D** Sceptic’s performance for scRNA-seq data of F123 (male) cells. **E** Sceptic’s performance for scRNA-seq data of F121-6 (female) cells. **F** Probability density function of Sceptic’s pseudotime for day 7 cells of scRNA-seq data.

To follow up on this observation, we next use Sceptic to infer pseudotime from scRNA-seq data generated from the same series of mESC samples [1]. For this data, Sceptic achieves an accuracy of 93.2% for male and 93.4% for female across five time points (Figure 5D–E). As in the ATAC-seq setting, we found for day 7, that female cells have a higher proportion of cells classified as earlier time points (12.4%; 62 out of 500 cells) than their male counterparts (2.5%; 28 out of 1102 cells) (Figure 5D–F), and similar proportions difference are observed for cells in day 11 (18.6% for female cells and 14.3% for male cells). This is consistent with our observation on scATAC-seq, but we found the transcript delay pattern is much stronger than the delay of chromatin accessibility.

### 3.5 Sceptic identifies methylation delay using co-assay data

We also applied our Sceptic model to a co-assay data, sc-GEM, a protocol that simultaneously measure pre-selected DNA methylation and gene expression markers in single cells via polymerase chain reaction (PCR) [5]. Cheow *et al*. performed sc-GEM on human fibroblasts undergoing induced pluripotent stem (iPS) cell reprogramming. This dataset contains 177 cells from five time points, BJ (day 0; fibroblasts established from skin taken from normal foreskin from a neonatal male), day 8, day 16, day 24 and iPS.

For this data set, Sceptic achieves 92.7% test set accuracy using gene expression data but only 63.8% using DNA methylation data (Figure 6). In particular, using methylation data, Sceptic can accurately distinguish among the later three time points well but struggles to distinguish between day 0 and day 8 cells. This observation suggests that there are enough transcriptomic changes within eight days of the reprogramming process to distinguish untreated cells and day 8 cells, but it takes longer than eight days to distinguish DNA methylation state changes. Sceptic’s results are consistent with a previous study, which observed that most of the DNA methylation changes in iPS reprogramming only occur after day 9 [12].

**Figure 6:**
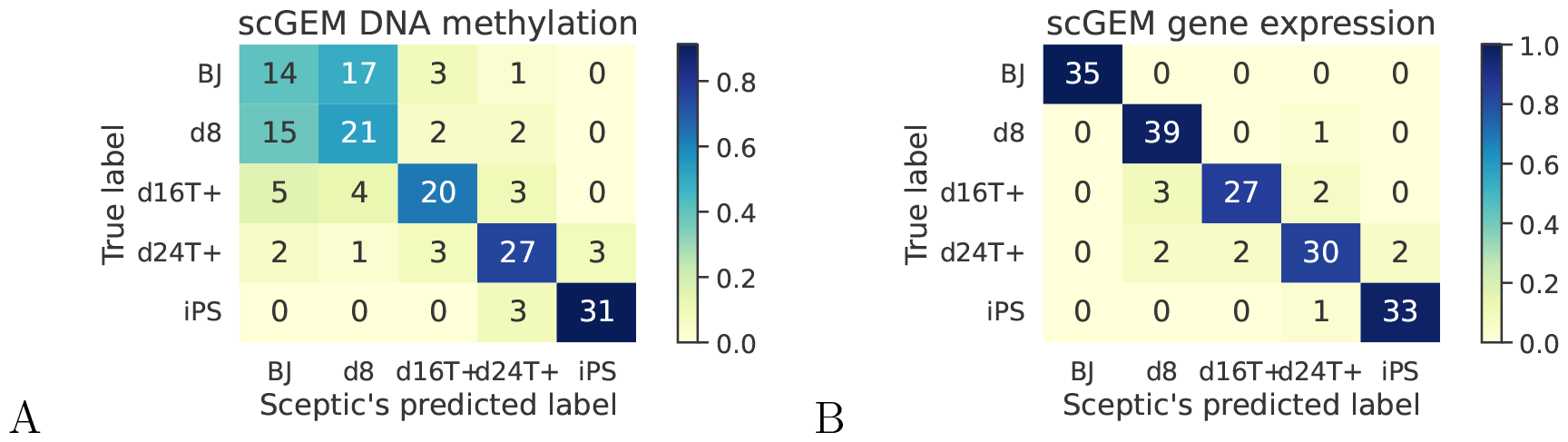
Sceptic captures DNA methylation delay. **A** Sceptic can also separate cells using DNA methylation data, albeit with some ambiguity between day 0 (BJ) and day 8. **B** Sceptic can accurately distinguish cells from 5 times points using gene expression data.

We explored the concordance between pseudotimes inferred from different modalities. To investigate this, we performed a two-way analysis of variance (ANOVA) to evaluate the influence of RNA-inferred pseudotime and time on methylation pseudotime. Notably, the analysis revealed a significant impact of RNA pseudotime on the variation in methylation pseudotime (P-value = 4.20e-11). This finding implies that the pseudotimes generated by two Sceptic models capture common underlying structures within the same cells, highlighting their shared aspects.

We also investigated Sceptic’s ability to work with single-cell Hi-C data, using allelic embryonic stem cell scHi-C data from a recent study [1]. Across five time points, Sceptic achieves 87.47% test set accuracy (Supplementary Section 2).

## 4 Discussion

In this work, we developed a method, Sceptic, to infer pseudotime for single-cell sequencing and imaging data using a supervised machine learning approach. Our experiments suggest that Sceptic offers improved accuracy relative to the previous supervised method, psupertime. Furthermore, to our knowledge, Sceptic is the first supervised computational approach that infers pseudotime using time-series single-cell ATAC-seq, single nucleus imaging data, and single-cell Hi-C data. Our extensive evaluation demonstrates that Sceptic effectively infers pseudotime for time-series sequencing and imaging data. Application of Sceptic to multiple single-cell RNA-sequencing data has shown uniformly improved performance compared with state-of-the-art methods. The extension to scATAC-seq and single nucleus imaging is straightforward and effective, and it successfully captures the cells in the same cell trajectory but from different time points and the cell heterogeneity within the same time points, providing a better time label for each cell.

There are several potential avenues for improvements to Sceptic. First, whereas psupertime’s linear model is relatively easy to interpret, Sceptic’s non-linear kernel functions make such interpretation more challenging. Intuitively, for more complex data structures like imaging, it is natural to use non-linear decision boundaries for high-dimensional classification problems. But it would still be interesting to find which genes are significantly important for Sceptic’s classifiers. Permutation tests could be an effective option when the number of genes and the number of cells are not too large.

Second, Sceptic could be made more scalable. In this work, we focused mainly on datasets of intermediate to small size. With the decreasing costs of sequencing, datasets containing millions of cells are becoming more common. Thus, future work could explore the scalability of the Sceptic model for application to such datasets. The current SVM implementation in Sceptic will likely be impractical beyond tens of thousands of cells. For large datasets, LinearSVC or SGD Classifier might be feasible. With enough cells to train, one could also take advantage of recently developed deep learning techniques to potentially further improve the classification performance. Note that a simple neural network did not work well for our mESC data, and we speculated that the main reason was an insufficient amount of training data.

Third, our current Sceptic model has been shown to work well for a simple cell differentiation trajectory or reprogramming process. However, one might consider how to compare pseudotimes across multiple concurrent lineages or cell trajectories.

Finally, when the sampling time points are very dense (say, sequencing every six hours) and the number of time points is large (say, greater than 40), then one might also consider formulating the pseudotime analysis problem directly as a regression problem. Along these lines, Calderon *et al*. adopted a neural network as a regression tool to infer pseudotime for *Drosophila* embryonic development [3]. Because their dataset was collected at overlapping time windows and we mainly focused on classification evaluation, we did not evaluate this dataset. It would be great to combine both the classification and regression nature of pseudotime analysis into a single loss function.

## References

[1] G. Bonora, V. Ramani, R. Singh, H. Fang, D. L. Jackson, S. Srivatsan, R. Qiu, C. Lee, C. Trapnell, J. Shendure, et al. Single-cell landscape of nuclear configuration and gene expression during stem cell differentiation and X inactivation. Genome Biology, 22(1):1–36, 2021.

[2] J. C. Caicedo, A. Goodman, K. W. Karhohs, B. A. Cimini, J. Ackerman, M. Haghighi, C. Heng, T. Becker, M. Doan, C. McQuin, et al. Nucleus segmentation across imaging experiments: the 2018 data science bowl. Nature methods, 16(12):1247–1253, 2019.

[3] D. Calderon, R. Blecher-Gonen, X. Huang, S. Secchia, J. Kentro, R. M. Daza, B. Martin, A. Dulja, C. Schaub, C. Trapnell, et al. The continuum of drosophila embryonic development at single-cell resolution. Science, 377(6606):eabn5800, 2022.

[4] J. Cao, M. Spielmann, X. Qiu, X. Huang, D. M. Ibrahim, A. J. Hill, F. Zhang, S. Mundlos, L. Christiansen, F. J. Steemers, C. Trapnell, and J. Shendure. The single-cell transcriptional landscape of mammalian organogenesis. Nature, 566(7745):496–502, 2019.

[5] L. Cheow, E. T. Courtois, Y. Tan, R. Viswanathan, Q. Xing, R. Tan, D. S. Tan, R. Robson, Y. Loh, S. R. Quake, et al. Single-cell multimodal profiling reveals cellular epigenetic heterogeneity. Nature methods, 13(10):833–836, 2016.

[6] M. Enge, H. E. Arda, M. Mignardi, J. Beausang, R. Bottino, S. K. Kim, and S. R. Quake. Single-cell analysis of human pancreas reveals transcriptional signatures of aging and somatic mutation patterns. Cell, 171(2):321–330, 2017.

[7] O. Genolet, A. A. Monaco, I. Dunkel, M. Boettcher, and E. G. Schulz. Identification of X-chromosomal genes that drive sex differences in embryonic stem cells through a hierarchical CRISPR screening approach. Genome Biology, 22(1):1–41, 2021.

[8] S. Gupta and R. Santoro. Regulation and roles of the nucleolus in embryonic stem cells: from ribosome biogenesis to genome organization. Stem Cell Reports, 15(6):1206–1219, 2020.

[9] K. He, G. Gkioxari, P. Dollar, and R. Girshick. Mask r-cnn. In Proceedings of the IEEE international conference on computer vision, pages 2961–2969, 2017.

[10] W. Macnair, R. Gupta, and M. Claassen. psupertime: supervised pseudotime analysis for time-series single-cell RNA-seq data. Bioinformatics, 38(Supplement 1):i290–i298, 2022.

[11] K. Pearson. Liii. on lines and planes of closest fit to systems of points in space. The London, Edinburgh, and Dublin philosophical magazine and journal of science, 2(11):559–572, 1901.

[12] J. M. Polo, E. Anderssen, R. M. Walsh, B. A. Schwarz, C. M. Nefzger, S. M. Lim, M. Borkent, E. Apostolou, S. Alaei, J. Cloutier, et al. A molecular roadmap of reprogramming somatic cells into iPS cells. Cell, 151(7):1617–1632, 2012.

[13] C. Qiu, J. Cao, B. K. Martin, T. Li, I. C. Welsh, S. Srivatsan, X. Huang, D. Calderon, W. S. Noble, C. M. Disteche, et al. Systematic reconstruction of cellular trajectories across mouse embryogenesis. Nature genetics, 54(3):328–341, 2022.

[14] X. Qiu, Q. Mao, Y. Tang, L. Wang, R. Chawla, H. A. Pliner, and C. Trapnell. Reversed graph embedding resolves complex single-cell trajectories. Nature methods, 14(10):979–982, 2017.

[15] W. Saelens, R. Cannoodt, H. Todorov, and Y. Saeys. A comparison of single-cell trajectory inference methods. Nature biotechnology, 37(5):547–554, 2019.

[16] E. G. Schulz, J. Meisig, T. Nakamura, I. Okamoto, A. Sieber, C. Picard, M. Borensztein, M. Saitou, N. Bluthgen, and E. Heard. The two active X chromosomes in female ESCs block exit from the pluripotent state by modulating the ESC signaling network. Cell stem cell, 14(2):203–216, 2014.

[17] M. Setty, V. Kiseliovas, J. Levine, A. Gayoso, L. Mazutis, and D. Pe’Er. Characterization of cell fate probabilities in single-cell data with Palantir. Nature biotechnology, 37(4):451–460, 2019.

[18] K. Street, D. Risso, R. B. Fletcher, D. Das, J. Ngai, N. Yosef, E. Purdom, and S. Dudoit. Slingshot: cell lineage and pseudotime inference for single-cell transcriptomics. BMC genomics, 19:1–16, 2018.

[19] T. N. Tran and G. D. Bader. Tempora: cell trajectory inference using time-series single-cell RNA sequencing data. PLoS computational biology, 16(9):e1008205, 2020.

[20] C. Trapnell, D. Cacchiarelli, J. Grimsby, P. Pokharel, S. Li, M. Morse, N. J. Lennon, K. J. Livak, T. S. Mikkelsen, and J. L. Rinn. The dynamics and regulators of cell fate decisions are revealed by pseudotemporal ordering of single cells. Nature Biotechnology, 32(4):381–386, 2014.

[21] B. Treutlein, Q. Y. Lee, J. G. Camp, M. Mall, W. Koh, S. A. M. Shariati, S. Sim, N. F. Neff, J. M. Skotheim, M. Wernig, et al. Dissecting direct reprogramming from fibroblast to neuron using single-cell RNA-seq. Nature, 534(7607):391–395, 2016.

[22] T. Wu, C. Lin, and R. C. Weng. Probability estimates for multi-class classification by pairwise coupling. In S. Thrun, L. K. Saul, and B. Scholkopf, editors, Advances in Neural Information Processing Systems, volume 16, pages 529–532, Cambridge, MA, 2003. MIT Press.

[23] Y. Yang, G. Li, H. Qian, K. C. Wilhelmsen, Y. Shen, and Y. Li. SMNN: batch effect correction for single-cell RNA-seq data via supervised mutual nearest neighbor detection. Briefings in bioinformatics, 22(3):bbaa097, 2021.

[24] Y. Yang, G. Li, Y. Xie, L. Wang, T. M. Lagler, Y. Yang, J. Liu, L. Qian, and Y. Li. iSMNN: batch effect correction for single-cell RNA-seq data via iterative supervised mutual nearest neighbor refinement. Briefings in Bioinformatics, 22(5):bbab122, 2021.

