## Supplementary material for "Pseudotime analysis for time-series single-cell sequencing and imaging data": Suppl

#### 1 Supplementary methods

##### 1.1 Imaging data

The image data are available upon request and in the process of submission to 4D Nucleome Consortium data portal `https://data.4dnucleome.org/li_sceptic_mESC_microscope`.

**Sample Collection** F121-6 mouse embryonic stem cells (ESCs) were grown and differentiated into embryoid bodies (EBs) as described in Bonora *et al.* [1]. Neural progenitor cells (NPCs) were derived from EBs at day 11 and collected at day 21. ESCs and EBs at days 3, 7, and 11, as well as NPCs were collected and frozen in 10% DMSO in 2–3 million cell aliquots.

**Immunostaining** Frozen cell aliquots were thawed at 37 C and resuspended in 1x DPBS. Approximately 250,000 cells were plated on a 24-well glass bottom, black-walled plates (CellVis, Mountain View, CA, P24-1.5H-N) via centrifugation at 300g for 5 mins. The cells were then fixed with 4% paraformaldehyde (Electron Microscopy Sciences Cat. No. 50-980-492) for 15 m at room temperature. After fixation, they were washed three times with 1x PBS and incubated in blocking buffer (3% BSA and 0.25% Tx100 in 1x PBS) for 1 hour at room temperature. They were then incubated with primary antibodies (Polyclonal nucleolin antibodies, 1:500, Invitrogen Cat. No.PA5-85972) diluted in blocking buffer for an hour at room temperature. The cells were washed three times with 1x PBS and incubated with secondary antibodies (Donkey anti rabbit Alexa 647, Invitrogen, Cat. No. A-31573) and DAPI diluted in blocking buffer for an hour at room temperature. They were then washed three times with 1x PBS and stored in 600 $\mu$L of 1x PBS.

**Image acquisition** Image acquisition was performed with an Automated Leica DMi8 inverted microscope with Adaptive Focus technology configured with a 40X 0.95 NA objective. The microscope was illuminated using a six-line Lumencor Spectra X Light Engine LED with Semrock multi-band dichroic filters (Spectra Services, Ontario, NY). The images were captured with a Zyla 4.2 sCMOS camera (Andor, Windsor, CT, USA) and the Piezo-driven stage (Okolab) was used to acquire z-stacks.

**Image segmentation** Nuclear segmentation was performed with a pre-trained Mask R-CNN model [2, 5] with a nuclear size cutoff of 900. Each nucleus was padded and cropped to a 128×128 pixel image.

**Image embeddings** For each single-nucleus image crop (128×128) of mESC dataset that we collected, we use pre-trained (on ImageNet [7]) Resnet34 [6](pytorch/vision:v0.10.0) from Pytorch [9] to generate a less-noisy vector-representation of the image. We generate 512-dim embeddings from the nucleolin-tagged image and 512-dim embeddings from the DAPI mask image. By concatenating them together, we generated a 1024-dim representation for each image crop.

#### 1.2 scATAC-seq data

We followed the epiScanpy pipeline to process the mESC scATAC-seq data [3]. Briefly, we first binarized the cell by peak matrix and identified the most variable peaks. The standard library size normalization and log normalization were performed. Finally, we used the top 50 principal components from normalized data as the input for our Sceptic model.

#### 1.3 scRNA-seq data

| Datasets | Type | Time Label | # time points | # cells | Accession |
|---|---|---|---|---|---|
| mESC [1] | sciRNA | Embryonic day | 5 | 4064 | GSE184554 |
| hESC [10] | scRNA | Embryonic day | 5 | 1529 | E-MTAB-3929 |
| Human germline [8] | scRNA | Age (weeks) | 12 | 992 | GSE86146 |
| Embryonic beta cells [11] | scRNA | Developmental stage | 7 | 575 | GSE87375 |
| Acinar cells [4] | scRNA | Donor age | 8 | 411 | GSE81547 |
| MEF to neurons [12] | scRNA | Days since induction | 5 | 315 | GSE67310 |

Table S1: **Time-series scRNA-seq datasets.**

We used six publicly available scRNA-seq data sets (Table S1). We downloaded the five scRNA-seq datasets that were previously analyzed by Manair *et al.* from the R package "psupplementary" (`https://github.com/wmacnair/psupplementary`). We also downloaded the mESC scRNA-seq and scATAC-seq datasets analyzed by Bonora *et al.* [1] from bitbucket repository (`https://bitbucket.org/noblelab/mouse-sci-omics/src/master/`). We downloaded the processed scGEM dataset from UnionCom's GitHub page `https://github.com/caokai1073/UnionCom/tree/master/scGEM`.

### 2 Sceptic works well for scHiC data

We also investigated whether Sceptic would generalize to single-cell HiC data. To do so, we collected allelic embryonic stem cell Tsixstop scHiC data from a recent study [1]. We downloaded the preprocessed contact decay profile for each chromosome and concatenated them across 20 chromosomes as the vector representation of each cell. To measure the classifier's performance, we randomly selected 83 cells from each time point, yielding 415 cells in total. Overall, Sceptic achieves 87.47% test set accuracy (Figure S1).

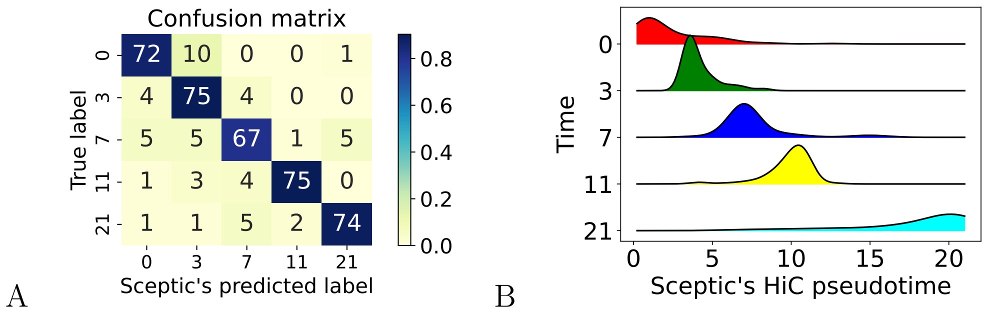


Figure S1: **Sceptic works well for scHiC data.** **A** Sceptic can also separate cells using allelic single-cell HiC data. **B** Ridge plot of Sceptic pseudotime, colored by the day of cells, on the Tsixstop data.

### References


[1] G. Bonora, V. Ramani, R. Singh, H. Fang, D. L. Jackson, S. Srivatsan, R. Qiu, C. Lee, C. Trapnell, J. Shendure, et al. Single-cell landscape of nuclear configuration and gene expression during stem cell differentiation and X inactivation. *Genome Biology*, 22(1):1–36, 2021.

[2] J. C. Caicedo, A. Goodman, K. W. Karhohs, B. A. Cimini, J. Ackerman, M. Haghighi, C. Heng, T. Becker, M. Doan, C. McQuin, et al. Nucleus segmentation across imaging experiments: the 2018 data science bowl. *Nature methods*, 16(12):1247–1253, 2019.

[3] A. Danese, M. L. Richter, K. Chaichoompu, D. S. Fischer, F. J. Theis, and M. Colomé-Tatché. Episcanpy: integrated single-cell epigenomic analysis. *Nature Communications*, 12(1):5228, 2021.

[4] M. Enge, H. E. Arda, M. Mignardi, J. Beausang, R. Bottino, S. K. Kim, and S. R. Quake. Single-cell analysis of human pancreas reveals transcriptional signatures of aging and somatic mutation patterns. *Cell*, 171(2):321–330, 2017.

[5] K. He, G. Gkioxari, P. Dollár, and R. Girshick. Mask r-cnn. In *Proceedings of the IEEE international conference on computer vision*, pages 2961–2969, 2017.

[6] K. He, X. Zhang, S. Ren, and J. Sun. Deep residual learning for image recognition. In *Proceedings of the IEEE conference on computer vision and pattern recognition*, pages 770–778, 2016.

[7] A. Krizhevsky, I. Sutskever, and G. E. Hinton. Imagenet classification with deep convolutional neural networks. In *Advances in neural information processing systems*, pages 1097–1105, 2012.

[8] L. Li, J. Dong, L. Yan, J. Yong, X. Liu, Y. Hu, X. Fan, X. Wu, H. Guo, X. Wang, et al. Single-cell rna-seq analysis maps development of human germline cells and gonadal niche interactions. *Cell stem cell*, 20(6):858–873, 2017.

[9] A. Paszke, S. Gross, F. Massa, A. Lerer, J. Bradbury, G. Chanan, T. Killeen, Z. Lin, N. Gimelshein, L. Antiga, A. Desmaison, A. Kopf, E. Yang, Z. DeVito, M. Raison, A. Tejani, S. Chilamkurthy, B. Steiner, L. Fang, J. Bai, and S. Chintala. Pytorch: An imperative style, high-performance deep learning library. In *Advances in Neural Information Processing Systems 32*, pages 8024–8035. Curran Associates, Inc., Vancouver, Canada, 2019.

[10] S. Petropoulos, D. Edsgärd, B. Reinius, Q. Deng, S. P. Panula, S. Codeluppi, A. P. Reyes, S. Linnarsson, R. Sandberg, and F. Lanner. Single-cell rna-seq reveals lineage and x chromosome dynamics in human preimplantation embryos. *Cell*, 165(4):1012–1026, 2016.

[11] W.-L. Qiu, Y.-W. Zhang, Y. Feng, L.-C. Li, L. Yang, and C.-R. Xu. Deciphering pancreatic islet $\beta$ cell and $\alpha$ cell maturation pathways and characteristic features at the single-cell level. *Cell metabolism*, 25(5):1194–1205, 2017.

[12] B. Treutlein, Q. Y. Lee, J. G. Camp, M. Mall, W. Koh, S. A. M. Shariati, S. Sim, N. F. Neff, J. M. Skotheim, M. Wernig, et al. Dissecting direct reprogramming from fibroblast to neuron using single-cell RNA-seq. *Nature*, 534(7607):391–395, 2016.
